# Role of the clathrin adaptor AP-1 in cell wall integrity and virulence factor secretion in the plant pathogen Botrytis cinerea

**DOI:** 10.1101/2024.12.19.629338

**Authors:** Glen Calvar, Adrien Hamandjian, Mélanie Crumière, Jean-William Dupuy, Christine Moriscot, Benoit Gallet, François-Xavier Gillet, Amélie de Vallée, Christine Rascle, Mathias Choquer, Christophe Bruel, Nathalie Poussereau

**Affiliations:** Microbiologie, Adaptation, Pathogénie, UMR5240, Univ Lyon, Université Lyon 1, CNRS, INSA, Bayer SAS, Lyon, France; Plateforme protéome, Centre de Génomique Fonctionnelle, Université de Bordeaux, Bordeaux, France; Integrated Structural Biology Grenoble (ISBG), Campus EPN / Bâtiment Institut de Biologie Structurale, Grenoble, France

**Keywords:** Adaptor protein 1, AP-1, growth, cell wall, secretion, clathrin, pathogenicity

## Abstract

In filamentous fungi, the secretory pathway plays a key role in polarized growth through the dlivery of vesicles carrying enzymes and components necessary to cell wall synthesis and fungal nutrition. In pathogenic species, it moreover supports the delivery of virulence factors. In metazoa and yeast, the formation of several secretory vesicles occurs at the Golgi apparatus and endosomes through a complex and partially characterized machinery involving clathrin and adaptor proteins such as AP-1. Data on the molecular mechanisms controlling this process are still scarce in filamentous fungi. Using a mutant under-expressing the β-subunit of the heterotetrameric AP-1 clathrin adaptor complex, we investigated the importance of intracellular vesicle trafficking in the phytopathogenic fungus *Botrytis cinerea*. The mutant exhibits pleiotropic developmental defects, an abnormality in cell wall integrity associated with a mislocalization of the plasma-membrane chitine synthase BcCHSIIIa. Moreover, the AP-1 mutant is affected in the secretion of hydrolytic enzymes and displays a severe defect in pathogenicity. This study confirms the importance of the AP1/clathrin machinery in the secretory pathway of *B. cinerea*.

## 1 Introduction

Efficient host-colonization by plant-pathogenic fungi relies on at least three main biological processes that are nutrient acquisition, hyphal growth, and secretion of virulence factors facilitating pathogen invasion. Moreover, hyphal growth is linked to the establishment of a functional cell wall that protects the fungus from external environmental stresses and particularly from the plant responses. All these processes rely on the production of secretory vesicles (SVs) and efficient vesicular trafficking.

The hyphal tip is considered the major site of exocytosis in filamentous fungi. SVs accumulate at the hyphal apex and fuse with the plasma membrane, delivering the lipids and the cell wall material required for polarized-cell elongation. Notably, SVs deliver fungal cell wall enzymes (FCWE) such as chitin synthases and the 1,3-β-glucan synthase and are essential for cell wall biogenesis (Verdín et al., 2019). In addition, SVs exocytosis is supposed to supportc the secretion of hydrolytic enzymes such as the xylanase XYNII, a Plant Cell Wall Degrading Enzyme (PCWDE) in *Trichoderma reesei* (Kurzątkowski et al., 1993), and a glucoamylase in *Aspergillus niger* (Wösten et al., 1991). This hypothesis is supported by several studies where the α-amylase *amyB* was localized in hyphal tip vesicles in several plant-pathogenic fungi (Hayakawa et al., 2011; Yang et al., 2021). In addition, virulence factors such as Cell Death Inducing Proteins (CDIP) are suspected to be secreted via SVs (Hayakawa et al., 2011; Yang et al., 2021). Thus, SVs could be considered a cornerstone of fungal virulence and yet very little is known about the mechanisms leading to their biogenesis.

Clathrin, a protein complex conserved in eukaryotes, serves as a structural scaffold in the biogenesis of both endocytic vesicles at the plasma membrane (PM) and SVs at the *trans*-Golgi network (TGN) and endosomes (Gurunathan et al., 2002; Borgonovo et al., 2006; Burgess et al., 2011; Robinson, 2015). It is made of heavy and light-chain subunits forming clathrin triskelions recruited to membranes by adaptor proteins (APs) and assembled to form a clathrin coat around nascent vesicles. AP-clathrin association likely contributes to the membrane deformation required for vesicle biogenesis (Makowski et al., 2020). After scission from the donor membrane, the clathrin/adaptor complex detaches from the vesicle during an uncoating process (Kirchhausen et al., 2014). Extensive studies have focused on clathrin-mediated endocytosis (CME) during the last 40 years (Robinson, 2015), but only very few described the role of clathrin in the production of secretory vesicles, particularly in filamentous fungi. A few years ago, clathrin was localized at the Late Golgi apparatus in *A. nidulans* (Schultzhaus et al., 2017; Martzoukou et al., 2018), consistent with its putative role in SVs biogenesis and evidence supports that clathrin plays a pivotal role in the secretion process of virulence factors in *B. cinerea* (Souibgui et al., 2021).

Clathrin does not directly bind cargo proteins as APs play a major role in sorting and recruiting transmembrane protein cargo. APs are heterotetramers of one small subunit σ (∼20 kDa), one medium subunit µ (∼50 kDa), and two large subunits β^1-5^ and either *α, γ, δ, ε, ζ* (∼100 kDa each). Subunits µ and σ sort transmembrane cargos by recognizing specific motifs located in their cytoplasmic tails and the β-subunit recruits coat proteins such as clathrin (Robinson, 2004). Up to five different APs (AP-1 to AP-5) have been found in animals and plants. Each AP is associated with vesicle formation at specific subcellular locations (Robinson, 2004; Hirst et al., 2013) and provides the specificity determining which cargo is selected. AP-1 considered the most indispensable and present in every eukaryote whose genome has been sequenced, acts in the formation of clathrin-coated vesicle trafficking between the TGN and early endosomes (EE) (Robinson, 2004; Robinson et al., 2024). AP-2 is implicated in clathrin-mediated endocytosis via its role in forming clathrin-endocytic vesicles at the plasma membrane. AP-3 could be involved in a clathrin-independent biogenesis of vesicles addressed to endosomes, lysosomes, or vacuoles at the TGN (Robinson, 2004; Nakatsu et al., 2014; Robinson, 2015). Finally, AP-4 and AP-5 do not interact with clathrin for vesicle biogenesis. AP-4 is implicated in the traffic between the TGN and specialized compartments, whereas AP-5 participates in trafficking between late endosomes and the TGN (Sanger et al., 2019). In filamentous fungi, only AP-1, AP-2, and AP-3 homolog complexes have been reported to be conserved (Martzoukou et al., 2017) and few functional studies have been described. In 2015, the absence of the AP-3 µ subunit was shown to be associated with a significant increase in lignocellulase secretion (Pei et al., 2015). In 2017, Martzoukou and his collaborators described that AP-2 was involved in a clathrin-independent endocytosis in *A. nidulans* (Martzoukou et al., 2017) and they showed in 2018 that AP-1 co-localizes with clathrin at the Golgi apparatus and is involved in SVs polar sorting (Martzoukou et al., 2018). More recently, the disruption of the AP-1 small subunit σ was shown to impact the physiology and the virulence of the phytopathogenic fungus *Fusarium graminearum* (Wu et al., 2023).

The characterization of a strain under-expressing the clathrin heavy chain we have previously constructed revealed that clathrin is critical to the secretion process of virulence factors in the phytopathogenic fungus *Botrytis cinerea* (Souibgui et al., 2021) suggesting a role for clathrin in the formation of SVs. To investigate this process further, we launched a functional study of AP-1, the only AP complex interacting with clathrin at the Golgi apparatus in filamentous fungi. The disruption of any AP-1 subunit has been reported to inactivate the function of the whole adaptor complex (Robinson, 2004, 2015). The gene encoding the β subunit of the AP-1 complex *(Bcap1b)* was chosen due to its critical role in clathrin recruitment. This study highlights the involvement of AP-1 in fungal elongation, cell wall integrity maintenance, hydrolytic enzyme secretion, and pathogenicity, emphasizing its critical role in both the saprophytic phase of the fungus and its necrotrophic strategy.

## 2 Material and methods

### 2.1 Fungal strains and culture conditions

*Botrytis cinerea* (teleomorph *Botryotinia fuckeliana* (de Bary Whetzel) strain B05.10 was used as a control and recipient for all genetic constructions.

Conidia were collected after 14 days of culture at 21°C under near-UV light (365 nm) on a modified Tanaka medium containing glucose (20 g.l^-1^), NaNO_3_ (2 g.l^-1^), KH_2_PO_4_ (2 g.l^-1^), MgSO_4_ (0.5 g.l^-1^), CaCl_2_ (0.1 g.l^-1^), saccharose (200 g.l^-1^), agar (15 g.l^-1^) and oligoelements (ZnSO_4_, CuSO_4_, H_3_BO_3_, MnSO_4_ and NaMoO_4_) as described in Tanaka medium.

Strains were cultivated on solid or in liquid MMII medium containing glucose (20 g.l^-1^), KH_2_PO_4_ (0.2 g.l^-1^), MgSO_4_·7H_2_O (0.1 g.l^-1^), KCl (0.1 g.l^-1^), FeSO_4_·H_2_O (0.002 g.l^-1^) and supplemented with NaNO_3_ (2 g.l^-1^) (inducing conditions) or glutamate (3 g.l^-1^) (repressive conditions).

Media were inoculated by depositing 10 µL of 10^6^ spores.ml^-1^ at the center of 90 mm-Petri dishes or by cultivating 600 spores per well in a 96-well TPP microplate containing liquid media. In the latter case, the optical density at 620 nm (OD620) was measured using SpectraMax Plus 384 Microplate reader at 0h and after 7 days of incubation in the dark at 21°C. For infection cushion (IC) formation, 600 spores per well were cultivated in liquid MMII medium supplemented with NaNO_3_ (2 g.l^-1^) in a 96-well glass base microplate and incubated for 3 days at 21°C. ICs were observed by inverted microscopy. Sclerotia formation was observed after 29 days of culture on modified Tanaka medium incubated at 21°C in darkness. To quantify conidiation, 7.5.10_5_ spores were inoculated on modified Tanaka medium and incubated at 21 °C under continuous near-UV light. Conidial production was analyzed by counting the conidia under a microscope after collecting them from 14-day-old cultures. Three independent biological replicates were at least performed.

### 2.2 Mutant construction in *Botrytis cinerea*

*BcΔBcap1b*, AP-1^cond^, *BcCHSIIIa-GFP*, AP-1^cond^*-BcCHSIIIa-GFP* strains were generated using a gene replacement strategy (Appendix S1, Fig. S1, Fig. S2, and Table S1). The constructions were cloned into *Agrobacterium tumefaciens* to transform *B*.*cinerea* using a protocol described by Rolland et al (2003).

### 2.3 RNA extraction and RT-qPCR

For *Bcap1b* gene (Bcin02g08180) expression analysis, conidia (10^5^ sp.ml^-1^) from the parental strain and the AP-1^cond^ mutant were first cultured in liquid minimal medium supplemented with NaNO_3_ (2 g.l^-1^) for four days (21°C, 110 rpm). Then, mycelia were filtered, split in two, and cultured for an hour in inducing (nitrate) or repressive (glutamate) conditions. RNA was extracted from 25 mg of freeze-dried mycelia using the RNeasy Midi Kit (Qiagen). RT-qPCR experiments were performed as previously described by Rascle et al. (2018) using ABI-7900 Applied Biosystems (Applied Biosystems). At least three independent biological replicates were performed. Relative quantification was used on the 2^-ΔΔCt^ method (Livak and Schmittgen, 2001). Genes encoding for actin (*BcactA*, Bcin16g02020), elongation factor (*Bcef1a*, Bcin09g05760) and pyruvate dehydrogenase (*Bcpda1*, Bcin16g01890) were used as normalization internal controls. Primers used for RT-qPCR are listed in Table S1. Primer pair P65/P66 is listed in (Table S1).

### 2.4 Transmission Electron Microscopy (TEM)

*Sample preparation-* B05.10, AP-1^cond^ and AP-1^cond^/C strains were grown in liquid MMII medium supplemented with 2 g.l^-1^ of NaNO_3_ (inducible condition) for four days at 21°C, 110 rpm. Mycelia were collected and fixed in 4% paraformaldehyde (PFA) and 0.4% glutaraldehyde (GA) in 0.1 PHEM buffer for 30 min at room temperature (RT) with gentle shaking. The buffer was discarded, and a second incubation (30 min, RT) was performed in 0.1M PHEM buffer containing 2% PFA and 0.3% GA. High-pressure freezing was performed as described in Mohamed et al. (Mohamed et al., 2021). Briefly, the mycelium was deposited on the 200-µm side of a 3-mm type A gold plate (Leica Microsystems) covered with the flat side of a 3-mm type B aluminum plate (Leica Microsystems) and vitrified using an HPM100 system (Leica Microsystems) as follows: after 40h at -90°C in acetone with 1% OsO_4_, the samples were slowly warmed to -60°C (2°/h), stored for 10h at -60°C, then the temperature was raised to -30°C (2°/h) and the samples stored for a further 10h at -30°C before an 1h incubation at 0°C and storage at -30°C until further processing (all these steps are performed automatically in a AFS2 instrument using a pre-defined program). Then, the samples were rinsed 4 times in pure acetone before being infiltrated with progressively increasing concentrations of resin (Epoxy embedding medium, Sigma) in acetone while increasing the temperature to -20°C. At the end, pure resin was added at RT and the samples were placed at 60°C for 2 days of polymerization.

*Transmission Electron Microscopy of fungal cells-* 70 nm thin sections were cut from resin-embedded samples from high-pressure-freezing, using a UC7 ultramicrotome (Leica Microsystems) and collected on 100 mesh Formvar carbon copper grids. The sections were post-stained 24h with 2% uranyl acette and lead citrate (5 minutes each). Samples were observed using a Tecnai G2 Spirit BioTwin microscope (Thermo Fischer Scientific) operating at 120 kV with an Orius SC100B CCD camera (Gatan).

### 2.5 Determination of susceptibility to osmotic and parietal stresses

Sensitivity to KCl, Nikkomycin Z and caspofungin was investigated by culture in MMII medium containing 2 g.l^-1^ of NaNO_3_ (inducible condition) placed in 96-well microplates. Solutions of KCl ranging from 0.005M to 2M were prepared directly in MMII medium. Each well was filled with 195 μL of KCl-supplemented MMII medium and inoculated with 5 μl of spores suspended in MMII at a concentration of 1.2 .10_6_ spores.ml^-1^. Stock solutions of Nikkomycin Z and caspofungin (Sigma) were prepared at 3 mg.ml^-1^ in DMSO before dilution in MMII medium to achieve final concentrations ranging from 0.04 μg.ml^-1^ to 400 μg.ml^-1^ for Nikkomycin Z and from 0.004 µg.ml^-1^ to 120 µg.ml^-1^for caspofungin. Each well was filled with 50 μl of drug-containing MMII and inoculated with 150 μl of spores suspended in MMII at a concentration of 4 .10_4_ spores.ml^-1^, ensuring a DMSO concentration below 1%. For all experiments, the optical density (OD) at 620nm was measured at 0h and 7 days post-inoculation to monitor fungal growth. Three biological replicates with 4 technical replicates were produced for each drug concentration.

### 2.6 Chitin quantification

A total of 5.10^6^ spores were cultured for three days at 21°C under 110 rpm in 50 ml of liquid MMII (inducible condition). Mycelia from liquid culture were harvested by filtration before deep-freezing in liquid nitrogen and lyophilization. Mycelia were ground using the *TissueLyser II* (Qiagen) and approximately 10-20 mg of ground mycelia were placed in three glass test tubes per strain. The weight of mycelium per tube was precisely measured. 5 ml of HCl 6 N were added to each tube before incubation at 100°C for six hours. The acidic reaction was neutralized by placing tubes on ice and by the addition of NaOH 5N until a pH of 6.0 to 7.0 was reached. The volume of each reaction mix was measured. Glucosamine was quantified using the modified Elson and Morgan method (Schloss, 1951). A volume of 100 μl of the hydrolysate was placed in a glass test tube before adding 650 μl of water and 250 μl of a solution containing Na^2^CO^3^ 1.25 N with 4% acetylacetone. Tubes were placed at 90°C for an hour and then on ice before adding 1.25 ml of absolute ethanol. After five minutes, 250 μl of Ehrlich’s reagent (1.6g DMAB in 30 ml HCl 12N and 30 ml ethanol) were added before incubation at room temperature in the dark for an hour. The absorbance at 520nm (OD_520_) was measured using a SpectraMax Plus 384 Microplate reader. The OD_520_ were converted to μg of D-glucosamine using a standard curve of 0 to 30µg of D-glucosamine.

### 2.7 Aniline blue fluorescence quantification

When excited at 395 nn, the aniline-blue/1,3-β-glucan complex emit fluorescence at 495 nm (Wood, 1984). Aniline blue fluorescence was investigated by culture in a 96-well microplate. Each well contained 600 spores suspended in 200 µl of MMII (inducible condition). At 7 days post-inoculation, the culture medium was discarded and 50 µL of 0.05% aniline blue in PBS pH 9.5 (Sigma) was added. After 60 min of incubation at 21°C, aniline blue fluorescence was measured using The Infinite® M1000 (Tecan).

### 2.8 General microscopy techniques and image acquisition

Confocal microscopy was performed with a Zeiss LSM850 confocal microscope (Oberkochen, Germany). 1,3-β-Glucan was stained with the addition of 0.05% aniline blue in PBS pH 9.5 (Sigma). Observations were performed an hour later using 405 nm laser excitation. For BcCHSIIIa-GFP localization, 72h-old growing hyphae in liquid MMII (inducible condition) were imaged (GFP excitation: 488 nm; emission 491 nm to 574 nm). FM4-64 10 µM (Sigma) was used as a lipophilic marker. Images were captured using the LSM510 software (Carl Zeiss) after 40 min of incubation with FM4-64. Images were stacked to produce Z-stacks with ImageJ (version 1.8.0).

### 2.9 Proteomic Analysis

A volume of 50 ml of liquid MMII (containing nitrate) was inoculated with spores (10^5^ sp.ml^-1^) and shaken (110 rpm) at 21°C in the dark for 96h. Mycelia were recovered by filtration, freeze-dried, and weighed. Culture filtrates were centrifuged at 14 000g for 15 minutes at +4°C, frozen in liquid nitrogen, and kept at −80°C before being used as samples in the proteomic analysis and the enzymatic reactions. Culture filtrates from four independent biological replicates were prepared as previously described for enzymatic analysis. After freeze-thawing overnight at 4°C, samples were centrifuged at 14 000g for an hour at 4°C to remove potential debris and extracellular polysaccharides. Proteins from culture supernatants were precipitated overnight at 4°C using trichloroacetic acid and sodium deoxycholate (15% and 0,05% final, respectively). Precipitated proteins were collected by centrifugation (14 000g, 4°C, 20 min), and washed two times with cold acetone. The pellet was air-dried for 30 minutes and resuspended in Laemli-DTT buffer (Tris 50mM, pH 6.8; EDTA 5 mM; Glycerol 10%; SDS 1%; bromothymol blue 0.02%; DTT 50 mM). Solubilized proteins were heated (98°C, 4 min) and stored at -20°C until proteomic preparation of the samples. Proteins (5 µg) were loaded onto a 10% acrylamide SDS-PAGE gel and visualized by Colloidal Blue staining. Migration was stopped when samples had just entered the resolving gel and the unresolved region of the gel was cut into only one segment. The steps of sample preparation and protein digestion by trypsin were performed as previously described (Rascle et al., 2018). NanoLC-MS/MS analysis were performed using an Ultimate 3000 RSLC Nano-UPHLC system (Thermo Scientific, USA) coupled to a nan-ospray Orbitrap Fusion™ Lumos™ Tribrid™ Mass Spectrometer Each peptide extract was loaded onto a 300 µm ID x 5 mm PepMap C18 pre-column at a flow rate of 10 µL/min. After a 3 min desalting step, peptides were separated on a 50 cm EasySpray column (75 µm ID, 2 µm C18 beads, 100 Å pore size,) with a 4-40% linear gradient of solvent B (0.1% formic acid in 80% ACN) in 91 min. The separation flow rate was set at 300 nL/min. The mass spectrometer operated in positive ion mode at a 1.9 kV needle voltage. Data were acquired using Xcalibur 4.4 software in a data-dependent mode. MS scans (m/z 375-1500) were recorded at a resolution of R = 120000 (@ m/z 200), a standard AGC target and an injection time in automatic mode, followed by a top speed duty cycle of up to 3 seconds for MS/MS acquisition. Precursor ions (2 to 7 charge states) were isolated in the quadrupole with a mass window of 1.6 Th and fragmented with HCD@28% normalized collision energy. MS/MS data was acquired in the Orbitrap cell with a resolution of R=30000 (@m/z 200), a standard AGC target and a maximum injection time in automatic mode. Selected precursors were excluded for 60 seconds. Protein identification and Label-Free Quantification (LFQ) were done in Proteome Discoverer 2.5. The MS Amanda 2.0, Sequest HT and Mascot 2.5 algorithms were used for protein identification in batch mode by searching against the ENSEMBL *B. cinerea* ASL83294v1 database (13749 entries, release 53). Two missed enzyme cleavages were allowed for trypsin. Mass tolerances in MS and MS/MS were set to 10 ppm and 0.02 Da. Oxidation (M) and acetylation (K) were searched as dynamic modifications and carbamidomethylation (C) as static modification. Peptide validation was performed using the Percolator algorithm and only “high confidence” peptides were retained, corresponding to a 1% false discovery rate at peptide level (Käll et al., 2007). Minora feature detector node (LFQ) was used along with the feature mapper and precursor ions quantifier. The quantification parameters were selected as follows: (1) Unique peptides, (2) Precursor abundance based on intensity, (3) Normalization mode: total peptide amount, (4) Protein abundance calculation: summed abundances, (5) Protein ratio calculation: pairwise ratio based, (6) Imputation mode: Low abundance resampling and (7) Hypothesis test: t-test (background based). Quantitative data were considered for master proteins, quantified by a minimum of 2 unique peptides, a fold changes above two, and an abundance ratio (for each biological replicate) seen 4 times with the same trend. The mass spectrometry proteomics data have been deposited to the ProteomeXchange Consortium via the PRIDE partner repository with the dataset identifier PXD037180.

### 2.10 Activities of secreted enzymes

Culture filtrates from four independent biological replicates were prepared as previously described for proteomic analysis. Enzymatic activities were performed as previously described by Souibgui et al., (2021). Briefly, protease activity was measured by incubating samples with 1% hemoglobin (Sigma), pH 3.5, and reactions were stopped with 25% trichloroacetic acid. After centrifugation, culture filtrates were mixed with NaOH 0.5 M and absorbance at 280 nm was recorded. Laccase activity was measured by incubating the samples and the ABTS substrate (Sigma) in 50 mM Na-acetate buffer, pH 4.0. Oxidation of ABTS was recorded at 405 nm (Molecular Devices Spectramax-485) during 45 min at 30°C. Xylanase and cellulase activities were recorded on a plate reader Tecan Infinite M1000 using the EnzChek-Ultra-xylanase assay kit (Thermofisher) and the cellulase assay kit (Abcam), respectively, according to manufacturers’ instructions. All activities were measured on three independent biological replicates. Pectin hydrolases and pectin-methyl-esterases activities were detected using pectoplates (Lionetti, 2015). Pectoplates were prepared with 0.1% (w/v) of apple pectin (Sigma 76282), 1% (w/v) agarose (Fisher), 12.5 mM citric acid and 50 mM Na_2_HPO_4_, pH 6.5. The gel was cast in Petri dishes and 60 ng of total protein in 40 μl from culture filtrates were deposited in 4mm diameter wells from culture filtrates Plates were incubated at 30°C for 24 h, and stained with 0.05% (w/v) Ruthenium red (Sigma) for 30 min. They were de-stained by several washes with ultrapure water and then photographed.

### 2.11 Pathogenicity assays

French bean leaves (*Phaseolus vulgaris* var Saxa), wounded apple fruit (*Malus domestica* cultivar Golden Delicious) and wounded tomato (*Solanum lycopersicum*), were inoculated with a conidia suspension (1,500 spores in 7.5µl of MMII medium supplemented with NaNO_3_ (2 g.l^-1^)) and incubated at 21°C under 80% relative humidity and dark-light (16 h/8 h) condition. Symptoms were scored up to 7 days post inoculation (dpi). Three independent biological replicates were at least assessed.

## 3 Results

### 3.1 *Bcap1b*, a gene involved in hyphal development

To investigate the role of the AP-1 complex in *B. cinerea*, we constructed a null mutant carrying the deletion of *Bcap1* (Bcin02g08180), the gene encoding the β-subunit of the complex (Fig. S1). However, heterokaryotic transformants carrying the deletion could not be purified into homokaryotic lines suggesting the essential role of this gene in the fungus. Given this result previously described in other organisms (Martzoukou et al., 2018), we generated a strain in which the *Bcap1b* gene was placed under the control of *pNiaD* (Fig. S2) a conditional nitrate reductase promoter from *B. cinerea* (Schumacher, 2012). A homokaryotic *pNiaD::Bcap1b* strain was obtained, hereafter named AP-1^cond^ mutant. This promoter induces the expression of the downstream gene in the presence of nitrate and represses it when other nitrogen sources are supplied (Schumacher, 2012). The quantification of *Bcap1b* expression confirmed its regulated expression in the mutant. The absence of nitrate in the medium (replaced by glutamate (3 g.l^-1^)) led to a 3.7-fold reduced *Bcap1b* expression compared with the parental strain but induced a severely retarded growth (Fig. 1A and 1B, Fig.S3). Microscopic observations of hyphae grown in the presence of glutamate revealed significant morphological alterations of hyphae, including swelling and hyperseptation (Fig. 1C). This severe growth defect compromised further phenotyping of the mutant strain in this condition.

**Figure 1.**
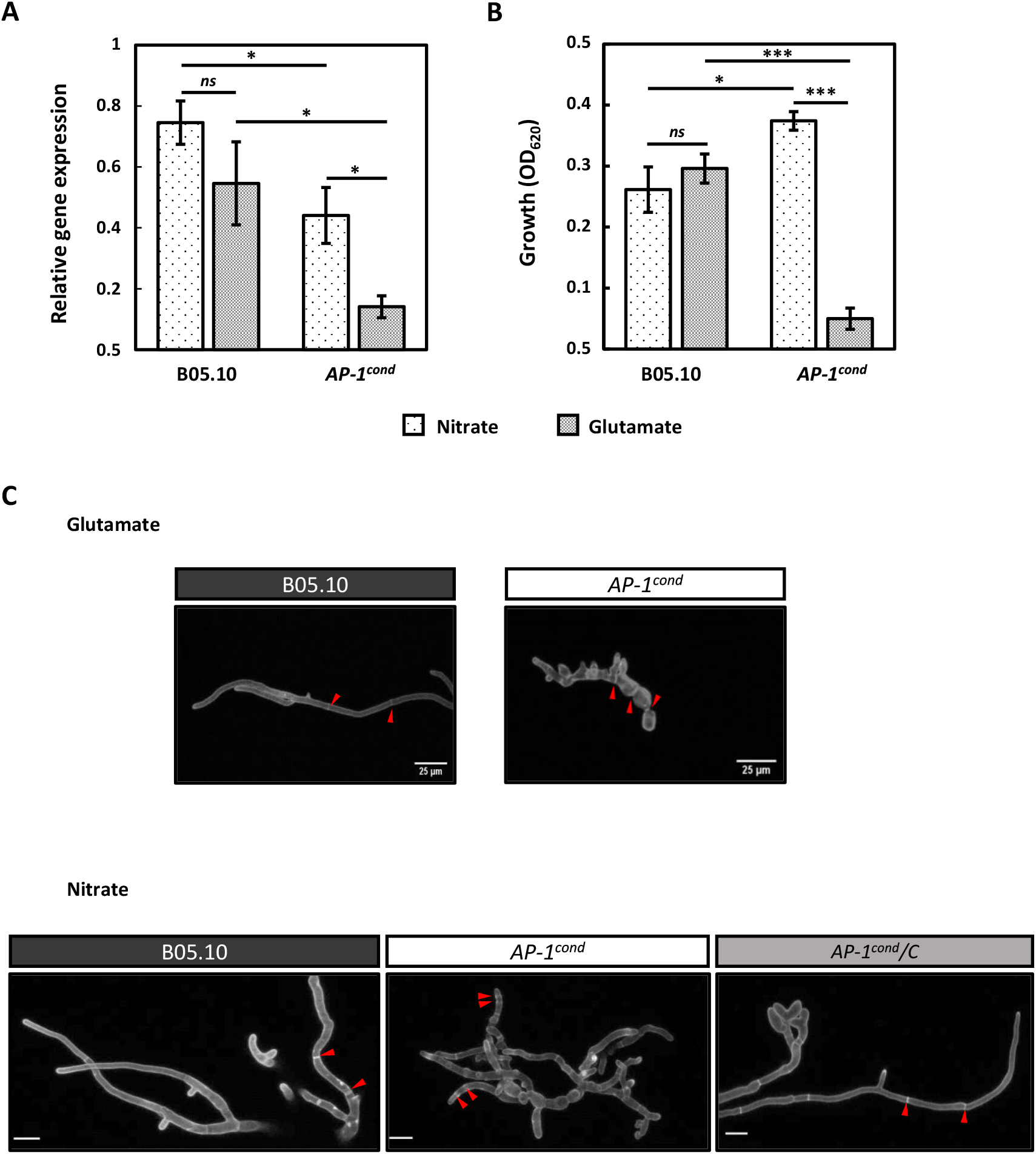
BcAp1b is involved in hyphal growth and morphology. **(A)** Comparative *Bcacp1*b expression patterns and growth of the B05.10 and AP-1^cond^ strains under inducible (nitrate) or repressive (glutamate) conditions. For *Bcacp1*b expression, mycelia were grown for 4 days in liquid minimal media and then transferred to media containing either nitrate (2 g.l^-1^) or glutamate (3 g.l^-1^) for 45 min. The actin -encoding gene *BcactA* was used as an internal control. **(B)** Growth quantification (OD_620_) of the B05.10 and AP-1^cond^ strains mycelia cultivated in liquid minimal medium in presence of nitrate or glutamate after 7 days of culture. All experiments were conducted with at least three biological replicates per conditions. Means with standard deviations are indicated and asterisks indicate a significant difference (Student’s *t*-test, * *p-*value < 0.05, ** *p-*value < 0.01, *** *p-*value < 0.001, *ns* non significant). **(C)** Confocal microscopy of hypha after 48h of growth in liquid minimal medium containing glutamate or nitrate. The cell wall was stained with aniline blue. Image acquisition was carried out using a 40x objective under 405 nm laser excitation. Septa are indicated with red arrows and hyphal swelling with white arrows. Scale bar = 10 µm.

In the presence of nitrate (2 g.l^-1^), *Bcap1b* expression was surprisingly reduced by 1.7-fold compared to the parental strain, and the growth rate of the mutant was not compromised as a slight increase (1.4-fold) of biomass was observed after 7 days of incubation (Fig. 1A and B, Fig.S3). When the mutant was inoculated on the same minimal nitrate medium but in solid conditions, an absence of radial growth was observed even after longer times of incubation (Fig.S4). Martzoukou et al., (2018) reported the same observation in the case of a conditional knock-down mutant of the encoding gene AP-1σ. Microscopy observations of hyphae grown in liquid conditions revealed that the AP-1^cond^ mutant displayed morphological defects with swelling (Fig. 1C), hyper-branching, and an abnormal septa distribution (Fig. 1C red arrows). Determination of the distance between septa revealed a 2.8-fold reduction in the mutant compared to the parental strain (Fig. S5). All these morphological alterations were no longer visible in the complemented strain. Additionally, hyphae of the AP-1^cond^ mutant were 60% wider compared to the parental strain, as illustrated by electron microscopy (Fig. 2A).

**Figure 2.**
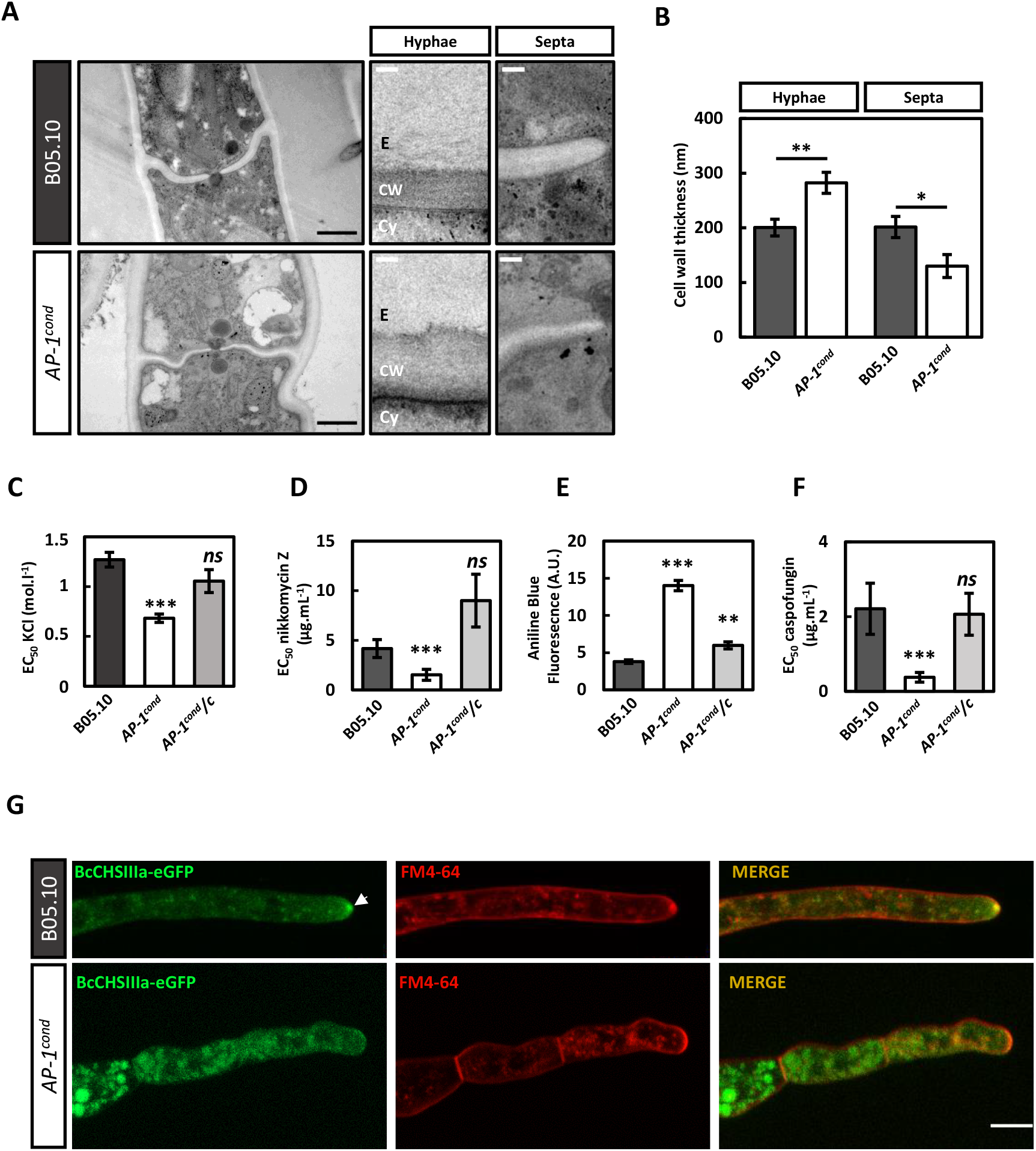
BcAp1b is involved in cell wall integrity maintenance. **(A)** Representative electron micrographs of *Botrytis cinerea* cultured in liquid minimal medium in the presence of nitrate showing the cell wall (CW), the cytoplasm and (Cy) and the extracellular medium (E) black scale bar = 500 nm; white scale bar = 75 nm. **(B)** Cell wall thickness of hyphae and septa. Fifteen electron micrographs were analysed, means with standard deviations are indicated (Student’s *t*-test, ** *p-*value < 0.01, * *p-*value < 0.05). **(C)** Sensitivity to KCl. Fungal growth was measured after 7 days of culture in minimal liquid media containing nitrate and supplemented with increasing concentrations of KCl. The relative half-maximal effective concentration (EC_50_) was calculated to estimate the susceptibility of the strains to KCl. Means with standard deviations are indicated (F-test, *** *p*-value < 0.001). **(D)** Chitin stress induced by Nikkomycin Z after 7 days of culture in minimal liquid media (inducible condition). Means with standard deviations are indicated (F-test, *** *p*-value < 0.001. **(E)** 1,3-β-glucan was quantified by measuring aniline-blue fluorescence after 7 days of culture in minimal liquid media (Inducible condition). Means with standard deviations are indicated (Student’s *t*-test, * *p-*value < 0.05, ** *p-*value < 0.01, *** *p-*value < 0.001, *ns* non significant) **(F)** 1,3-β-glucan stress induced by Caspofungin after 7 days of culture in minimal liquid media (inducible condition). Means with standard deviations are indicated (F-test, *** *p*-value < 0.0005). **(G)** Confocal micrographs of chitin synthase IIIa in the parental strain and *AP-1*^*cond*^ mutant. In B05.10, BcCHSIIIa-GFP localizes polarly at the fungal apex while in *AP-1*^*cond*^ mutant, BcCHSIIIa-GFP is mainly localized in small puncta in the subapical region. The lipophilic marker FM4-64 was used to label the plasma-membrane and plasma-membrane derived organelles. Images are maximum intensity projections from Z-stack .(Image J version 18).

### 3.2 *Bcap1b* contributes to cell wall integrity maintenance

Electron microscopy images revealed the importance of *Bcap1b* in cell wall biogenesis. Compared to the parental strain, the cell wall along the hyphae of the mutant strain was significantly 35% thicker. In contrast, the mutant septa were 27% thinner than those of the parental strain (Fig. 2A and 2B). These observations suggested that the AP-1^cond^ mutant may exhibit alterations in the composition and the properties of the cell wall. To investigate a potential difference in the cell wall integrity, the effect of increasing potassium chloride (KCl) concentrations on fungal growth was monitored in both parental and mutant strains. Using the relative half-maximal effective concentration (EC_50_) as an indicator, the collected data showed that the AP-1^cond^ mutant was highly sensitive to KCl, with a 1.9-fold reduction in EC_50_ compared to the parental and complemented strains (Fig. 2C). This higher sensitivity to osmotic stress in the AP-1^cond^ mutant suggested a modification of the cell wall composition. In the presence of nikkomycin-Z, a chitin synthase inhibitor, the AP-1^cond^ strain exhibited a 2.7-fold decrease in EC_50_ compared to the parental strain. (Fig. 2D). This increased susceptibility to nikkomycin-Z may indicate a potential alteration of the chitin synthesis. Supporting this hypothesis, quantification of glucosamine, a derivate of the chitin monomer N-acetyl-glucosamine revealed a 36% reduction in glucosamine content in the AP-1^cond^ mutant compared with the parental strain (Fig. S6). To investigate further, the localization of the class IIIa chitin synthase BcCHSIIIa was examined in both parental and mutant strains (Fig. S7). Using confocal microscopy, fine fluorescent particles marking the hyphal tip were observed in the parental strain producing BcCHSIIIa-GFP. In the AP-1^cond^ mutant, this signal was no longer apparent in the apical region but localized in larger puncta scattered throughout the cells (Fig. 2G). Thus, BcAp1b is required for the proper localization of BcCHSIIIa-GFP. When aniline blue was added to mycelia, the fluorescence emitted by the specifically stained 1,3-β-glucan (Wood et al, 1984) was 3.5 times higher in the AP-1^cond^ mutant when compared to the parental strain, suggesting an accumulation of this polysaccharide in the mutant cell wall (Fig. 2E). In connection with this, a strong sensitivity of the mutant strain to caspofungin, a 1,3-β-glucan synthase inhibitor was observed. The EC50 for caspofungin was 5.8 times lower in the AP-1^cond^ mutant than in the parental strain, indicating a significant increase in sensitivity to the drug (Fig. 2F). Altogether, these results demonstrate the critical role of *Bcap1b* in maintaining fungal cell wall integrity.

### 3.3 Bcap1b is involved in fungal differentiation programs

The parental and mutant strains were cultured on a solid modified Tanaka medium for 14 days to evaluate macroconidia production. Although radial growth was altered, the AP-1^cond^ mutant differentiated conidiophores and conidia with a phenotype similar to the parental strain and no difference in conidium germination was observed between the parental and the mutant strains. However, an 8-fold reduction in the number of conidia produced by the AP-1^cond^ mutant was measured in comparison with the parental strain indicating a strong impairment of asexual reproduction capacities (Fig. 3A). Longer incubation of the cultures confirmed that this defect was quantitative and not due to a delay in conidial differentiation. The sclerotium differentiation was also investigated on solid modified Tanaka medium but the plates were incubated in darkness for 29 days. The AP-1^cond^ mutant formed irregular-edged colonies and failed to produce sclerotia even after longer incubation (Fig. 3B). In the complemented strain, the aspect of the reproductive mycelium was not completely restored and a minor defect in sclerotia differentiation was noticed as the number of these structures was slightly reduced in comparison with the parental strain.

**Figure 3.**
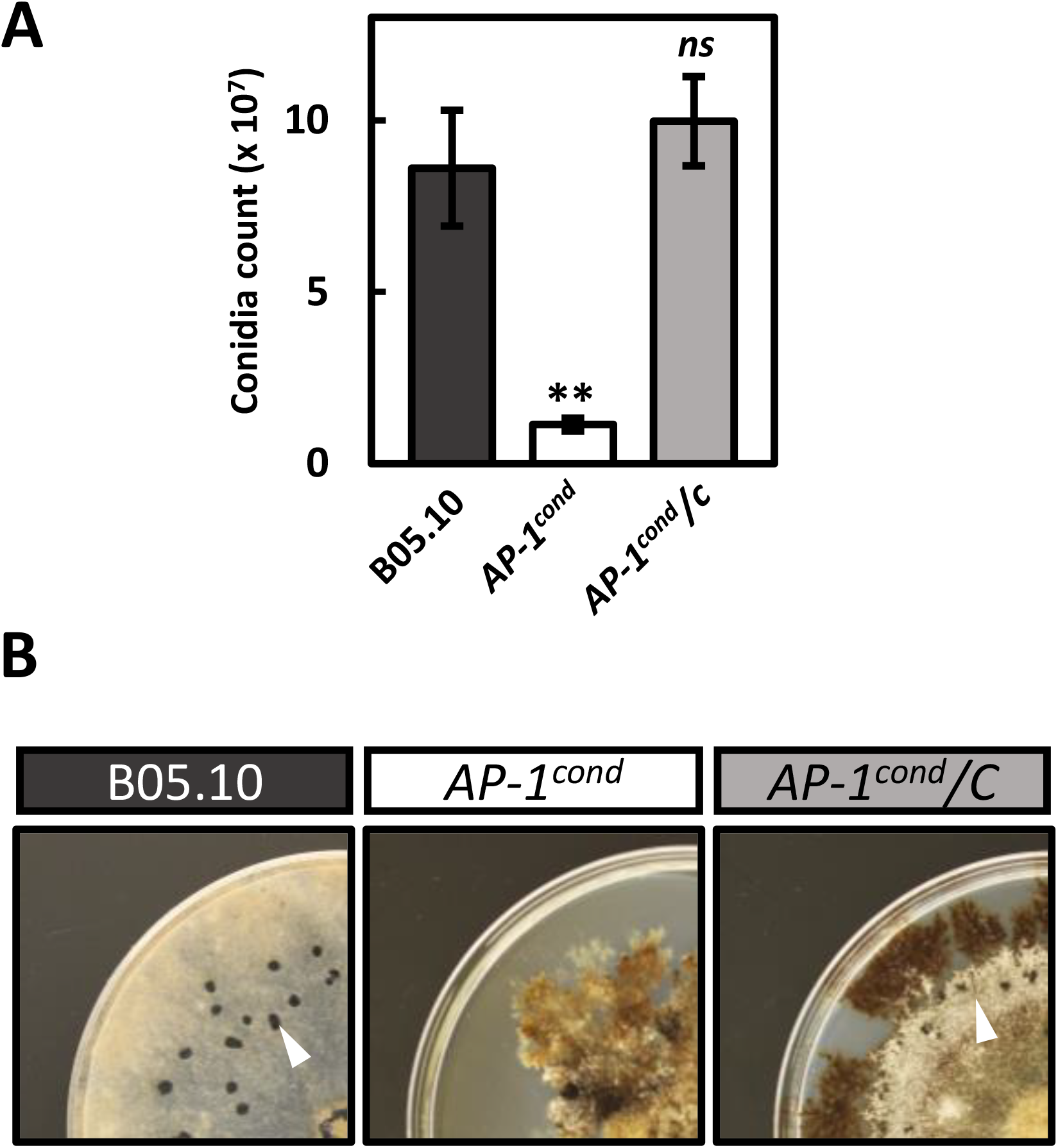
The *AP-1*^*cond*^ mutant shows defects in differentiation programs. **(A)** Conidiation was quantified from 14-day-old cultures grown on modified Tanaka medium (inducible condition) under near-UV light conditions. Spores were collected and counted under a microscope. Data were collected from three biological replicates. Means with standard deviations are indicated. (Student’s *t*-test, \*\**p*-value < 0.01, *ns* non-significant). **(B)** Sclerotium formation was observed after 29 days of culture on solid Tanaka medium maintained in the dark. White arrows indicate sclerotia.

### 3.4 *Bcap1b* is involved in protein secretion

In a previous study, we highlighted the important role of clathrin in the secretion process of virulence factors and hydrolytic enzymes in *B. cinerea* (Souibgui et al., 2021). This work suggested that clathrin could be involved in the biogenesis of secretory vesicles in plant-pathogenic fungi. As AP-1 is known to act together with clathrin in the biogenesis process of secretory granules in the fruit fly *D. melanogaster* (Burgess et al., 2011), we investigated the role of AP-1 in protein secretion in *B. cinerea*. For this purpose, the AP-1^cond^ and parental strains were cultured in a liquid MMII medium containing nitrate (2 g.l^-1^) to perform a comparative shotgun proteomic analysis of the secreted proteins. The quantification of secreted proteins, normalized to the produced biomass revealed that the AP-1^cond^ mutant accumulated more total extracellular proteins than the parental strain in its culture medium, suggesting an alteration of the secretion process (1.8-fold. Fig. S8). In 4 biological replicates, 343 proteins identified with a minimum of 2 unique and specific peptides were detected in the filtrate of both strains. Based on the presence of a signal peptide and the absence of a transmembrane domain, 185 proteins (54%) (Table S2) were predicted as secreted by the conventional secretory pathway. Surprisingly, 74% of these proteins (137 proteins) were down-accumulated in the mutant exoproteome when compared to the parental strain (Fig. 4A, Table S2) suggesting a crucial role of BcAp1b in the conventional secretory pathway. Functional classification of these differentially accumulated proteins highlighted 10 categories impacted in the AP-1^cond^ mutant (Fig. 4A). Protein degradation represented the most altered functional category, with 21 proteases down-accumulated and only 4 up-accumulated. Five members of the aspartic protease family were found down-accumulated and the aspartic protease BcAP8 (Bcin12p02040.1) was one of the most down-accumulated proteins (500-fold) in the mutant exoproteome. The plant cell wall degrading enzymes (PCWDEs) category was also highly altered, with 30 proteins down-accumulated, among which 14 pectin-associated enzymes including the endo polygalacturonase BcPG2 (Bcin14p00610.1 - 48-fold down), the endo arabinanase BcARA1 (Bcin02p07700.1 – 37-fold down) identified as virulence factors (Kars et al., 2005; Nafisi et al, 2014). On the other side, the polygalacturonase BcPGA1 (Bcin14p00850.1), an important factor in the infectious process (Have et al., 1998) was up-accumulated in the mutant (33-fold up). Finally, hemicellulose, hemicellulose-pectin (HP) and cellulose-related PCWDEs were also strongly down-accumulated. Among them, the endo-β 1-glucanase Cel5A (Bcin03p0400) was 200-fold less accumulated in the mutant. ‘oxidoreduction’ and ‘lipid metabolism’ were the two additional categories much altered by the mutation. All 13 proteins related to lipid metabolism were down-accumulated, and 14 of the 17 proteins related to oxidoreduction were also down-accumulated, with a strong negative ratio for the laccases BcLCC2 (Bcin14p02510.1 - 500-fold down) and BcLCC8 (Bcin01p00800.1–25-fold down). Finally, the fungal cell wall-related enzymes (FCWE) category was also concerned by the mutation with eight members up-accumulated (up to 16.4-fold) and 14 members down-accumulated (up to 167-fold) including the chitin-binding protein BcLysM1(Bcin02p05630 – 166-fold down), an effector that protects hyphae from chitinases (Crumière et al., 2024), and the transglycosylase BcCRH1(Bcin01p06010.1–22-fold down), considered as a cell death-inducing protein (Bi et al., 2021).

**Figure 4.**
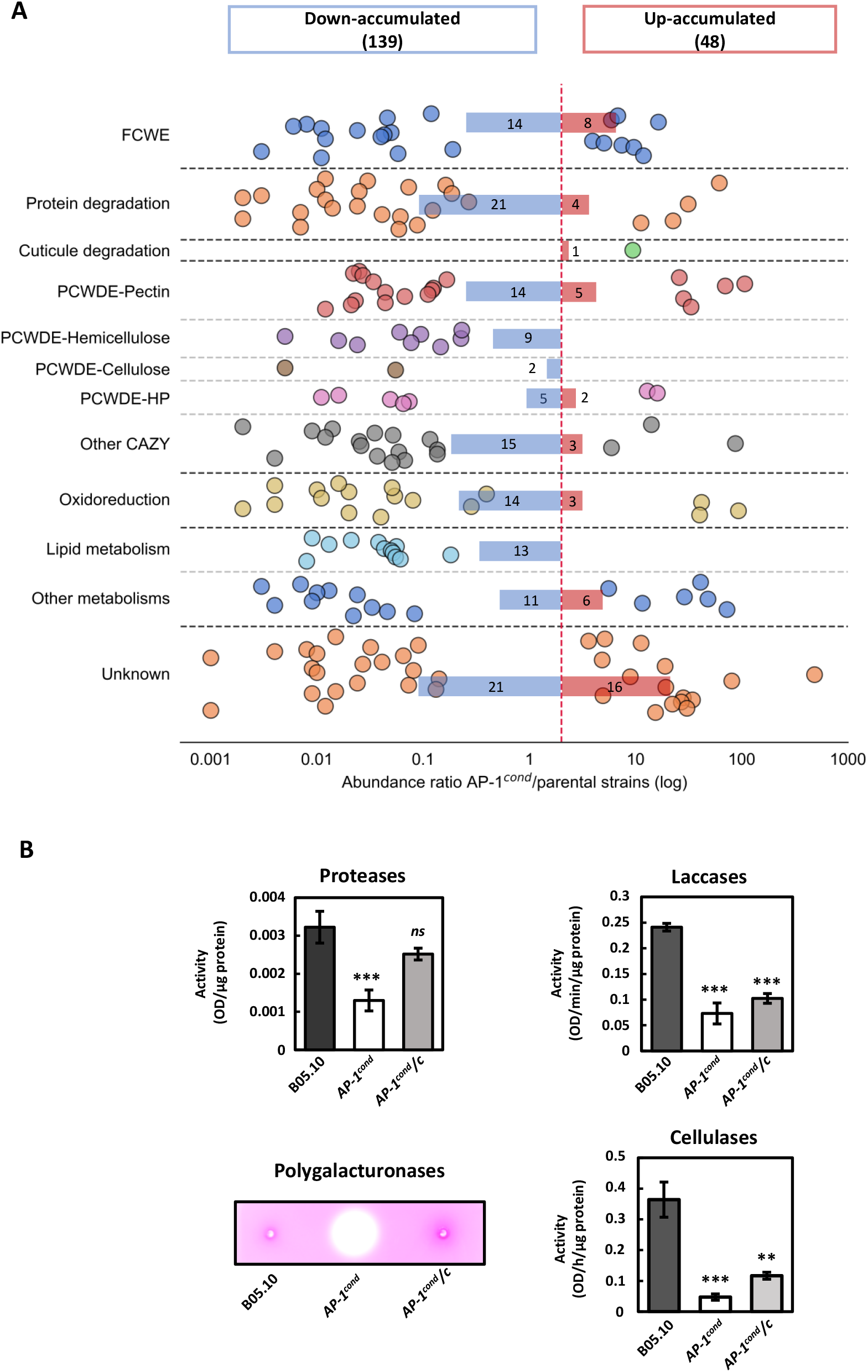
BcAp1b is involved in the secretion of hydrolytic enzymes. **(A)** Functional category classification of predicted secreted that are up- or down-accumulated in *AP-1*^*cond*^ mutant compared to the parental strain B05.10 (fold change > 2 ; see Supplementary Table 2 for details). For each protein (coloured circles), the abundance ratio between the AP-1cond mutant and the parental strain is shown. Analyses were performed after 4 days of growth in liquid minimal medium containing nitrate (four independent biological experiments). CAZy, carbohydrate active enzymes; PCWDE, plant cell wall degrading enzymes; FCWE, fungal cell wall enzymes; HP, Hemicellulose-Pectin. **(B)** Enzymatic activities quantified in the supernatants of liquid cultures in minimal liquid medium (inducible condition) at 4 days post-inoculation for the B05.10, *AP-1*^*cond*^ and *AP-1*^*cond*^*/C* strains. Three independent biological replicates were performed. Means with standard deviations are indicated (Student’s *t*-test, * *p-* value < 0,05**, *p-*value < 0.01, *** *p-*value < 0.001). Visualized detection of polygalacturonases on pectoplate, inoculated with four-day-old culture supernatants from the B05.10, *AP-1*^*cond*^ and *AP-1*^*cond*^*/C* strains. The photographs are representative of four independent biological replicates.

In connection with the functional categories highlighted by the proteomic analysis, proteases, laccases, and cellulases activity assays were quantified in the culture media used for proteomic analysis (Fig. 4B). When compared to the parental strain, protease, laccase and cellulase activities were reduced to 2.5, 3, and 7.4-fold respectively. While these results were in agreement with the proteomic data, a pectoplate assay (Lionetti, 2015) revealed, a higher pectin hydrolase activity, in contrast to the parental strain (Fig. 4B). All these results indicate that *Bcap1b* is important for the secretion of some enzymes involved in plant degradation or fungal cell wall synthesis.

### 3.6 *Bcap1b* is critical for pathogenicity

To evaluate the impact of AP-1 on the virulence of *B*.*cinerea*, french bean leaves were confronted with a conidial suspension of the parental, AP-1^cond^ and AP-1^cond^ /C strains. At 7 days post-inoculation, leaves infected by the mutant revealed very limited primary lesions in contrast to the parental and complemented strains that showed extensive tissue maceration (Fig. 5A). Identical results were obtained on tomato and apple fruits (Fig. 5B and C). Whatever the tested hosts, no evolution in symptom maceration was noticed with the mutant. To complete these observations and to investigate the early steps of infection, conidial suspensions were deposited onto the onion epidermis and onto glass slides to test and visualize the capacity of the fungus to penetrate plant cells and differentiate infection cushions, multicellular appressoria involved in plant penetration. To complete these observations, conidial suspensions of the mutant were deposited onto the onion epidermis and their ability to penetrate plant cells was visualized While hyphae of the parental and complemented strains penetrated the plant cells, the AP-1^cond^ mutant grew on the onion epidermis and failed to penetrate even after a longer time of incubation (Fig. 5D).

**Figure 5.**
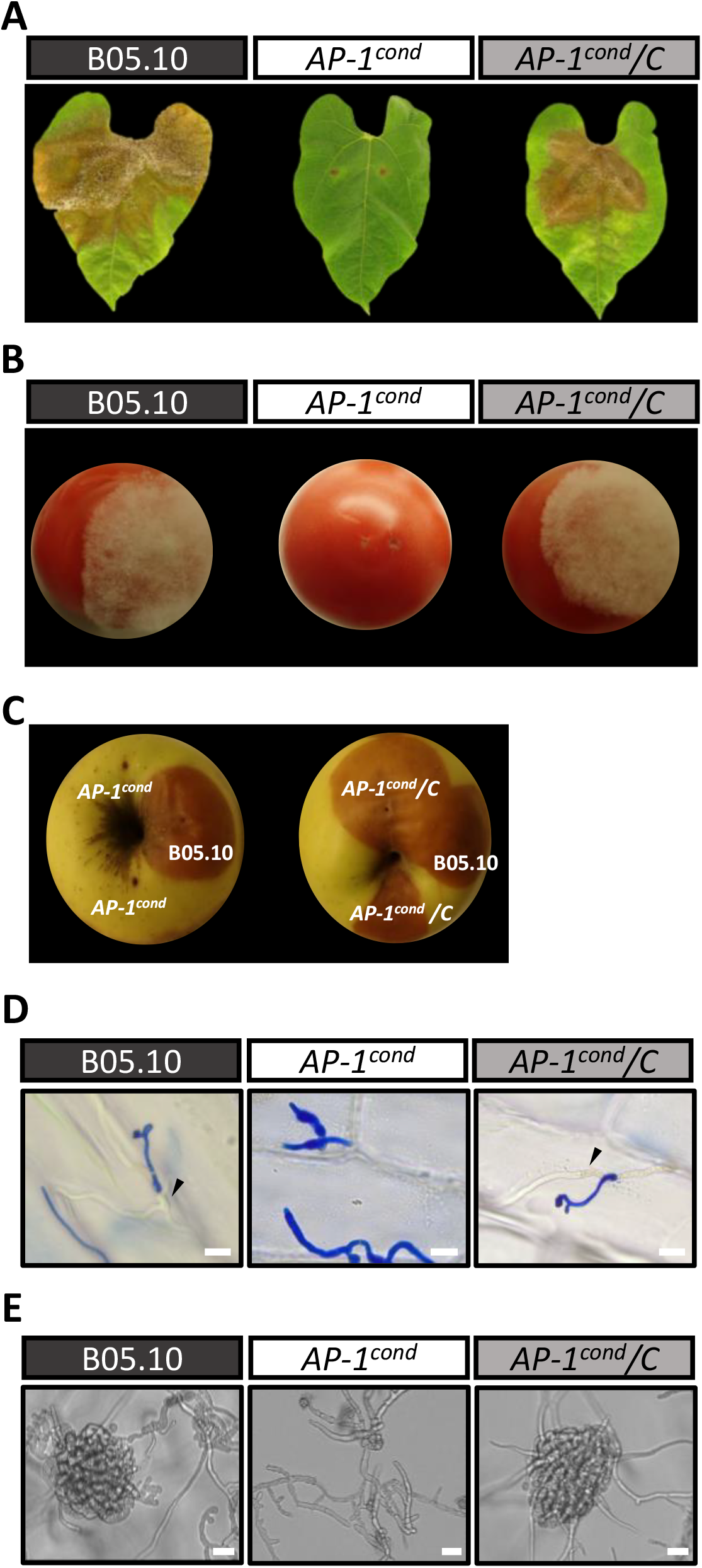
BcAp1b is important for pathogenicity. Virulence of B05.10, *AP-1*^*cond*^ and *AP-1*^*cond*^*/C* strains on French bean leaves **(A)**, tomatoes **(B)** and apple fruits **(C)**, 14 days old conidia were used for inoculation. Pictures were taken seven days post-inoculation. **(D)** Infection cushions were observed by inverted microscopy 3 days after inoculation in inducible liquid media on glass surfaces (scale bar 25 µm). **(E)** Penetration on onion epidermis: pictures were taken after 21h inoculation with conidia of B05.10, *AP-1*^*cond*^ and *AP-1*^*cond*^*/C*. Penetration sites are indicated with black arrows (scale bar 20 µm).

The capacity of differentiate infection cushions, multicellular appressoria involved in plant penetration was determined with spores deposited on glass cover slides. While hyphae of the parental and complemented strains produced infection cushions, the AP-1^cond^ mutant failed in this differentiation step (Fig. 5E).

## 4 Discussion

For several decades, the highly conserved clathrin adaptor AP-1 has been known for its role in protein trafficking between the TGN and the endosome. It is now well established that AP-1 is also involved in polarized protein trafficking. This role has been described not only in mammalian epithelial cells but also in plant and filamentous fungal cells (For review, Duncan 2022).

Here, we investigated the role of AP-1 and highlighted its importance in the fungal biology and the infectious strategy of the necrotrophic fungus *B. cinerea*.

### AP-1 is indispensable for fungal elongation and cell wall synthesis

The AP-1 complex is composed of 4 subunits and the disruption of any subunit has been reported to inactivate the function of the whole adaptor complex (Robinson, 2004, 2015). Therefore, we focused on the β-subunit that recruits clathrin during clathrin coated vesicle formation. Attempts to create a *Bcap1β* deletion mutant did not lead to homokaryotic strains. This is consistent with the report that AP-1 complex is essential in many organisms, including the filamentous fungus *A. nidulans* (Martzoukou et al., 2018). Hence, we performed a conditional genetic approach using the nitrate reductase promoter, a regulated promoter frequently used in *B. cinerea* (Schumacher, 2012; Marchegiani et al., 2015). In *A. nidulans*, Martzoukou et al., (2018), reported a growth alteration of a conditional knock-down mutant of AP-1σ encoding gene. Similar results were obtained in the conditional AP-1β (AP-1^cond^) mutant of *B. cinerea*, where polar growth was strongly impaired, exhibiting wider, shorter hyphae with more numerous septa. Similar observations were also reported in *A. niger* when *aplD* (AP-1*γ* encoding gene) was downregulated (Cairns et al., 2019). Interestingly, in the yeasts *S. cerevisiae* and *S. pombe*, which do not exhibit polar growth, AP-1 is not essential (Stepp et al., 1995; Huang et al., 1999; Valdivia et al., 2002; Ma et al., 2009; Yu et al., 2012; Arcones et al., 2016). In plants, AP-1 is also not essential but is required for root growth in *A. thaliana* (Park et al., 2013). These elements seem to confirm the hypothesis formulated by Martzoukou *et al*. (Martzoukou et al., 2018), who proposed that AP-1 is essential in filamentous fungi because filamentous growth depends on polarized vesicle trafficking and, in particular, apical sorting of polar cargoes, including those required for plasma membrane and cell wall biosynthesis.

The abnormal localisation of the class III chitin synthase BcCHSIIIa-GFP in the AP-1^cond^ mutant of *B*.*cinerea* supports this hypothesis. Indeed, the detection of the tagged protein in larger puncta scattered throughout the cells rather than concentrated in the apical region, indicates a mislocalization of this plasma membrane protein. This observation suggests a disruption in the trafficking process of this cell wall synthase to the hyphal tip. Interestingly, the chitin synthase ChsB (BcCHSIIIa ortholog) also mis-localizes in AP-1 mutants of *A. nidulans* (Martzoukou et al., 2018). Abnormal addressing of the class III chitin synthase likely could explain the alteration of the cell wall content, integrity and beyond the aberrant morphological features observed in the AP1 mutant. BcCHSIII is known as a major chitin synthase in *B cinerea* (Soulier et al., 2006) and we can question whether the trafficking of the other chitin synthase is dependent on AP-1 or not. Observation under electron microscopy revealed a cell-wall significantly thicker in the AP-1^cond^ mutant suggesting an over-accumulation of 1,3-β-glucan that may compensate for the chitin defect. This hypothesis is further supported by the hypersensitivity of the AP-1^cond^ mutant to the 1,3-β-glucan synthase inhibitor caspofungin. Thus, when this mutant is exposed to caspofungin, the production of 1,3-β-glucan is impaired, and the mutant fails to compensate for the chitin defect and becomes more sensitive to the drug. Thus, contrary to the chitin synthase CHSIIIa, the traffic of the glucan synthase BcFKS1 towards the hyphal tip might not depend on AP-1 in *B. cinerea*. Interestingly, in the fission yeast *S. pombe*, the glucan synthase BGS1 is a cargo of AP-1 (Yu et al., 2012). Further work is therefore required to determine whether or not the transport of the glucan synthase FKS1 is AP-1-dependent in filamentous fungi.

### AP-1 is involved in the secretion process of *B. cinerea*

As late secretory processes rely on AP-1 in many eukaryotes, we investigated the impact of the reduced expression of *Bcap1b* on the secretion capacity of *B. cinerea*. To our surprise, total protein quantification showed a higher amount of secreted proteins in the culture medium of the AP-1^cond^ mutant compared to the parental strain. In *A. niger*, Cairns et al., (2019), also reported a putative increase in protein secretion when *aplD* (AP-1*γ* subunit) was downregulated. They hypothesized that the possible hypersecretion phenotype would be the result of hyperbranching. We also observed this phenotype in AP-1^cond^ mutant. The hypersecretion phenotype might also be linked to the hyper-septation observed in the AP-1^cond^ mutant. Indeed, septa are known as exocytic sites for the delivery of hydrolytic enzymes such as the amylase *amyB* in *A. oryzae* and *F. odoratissimum* (Hayakawa et al., 2011; Yang et al., 2021). The increased number of septa observed in the AP-1^cond^ mutant may result in an increased number of exocytosis sites. The localization of the exocyst complex components in the AP-1^cond^ mutant could further address this hypothesis.

A quantitative proteomics approach revealed that AP-1 is involved in the secretion process of PCWDEs, proteases and oxidoreduction-related proteins. Interestingly, the secretion defect observed in the AP-1^cond^ mutant displays a similar signature to *B. cinerea* clathrin mutants (Souibgui et al., 2021) (Table S2). Indeed, protease, cellulase and laccase activities are also reduced in the clathrin mutants. Our results suggest that a functional AP-1/clathrin machinery is required for the secretion process of several enzymes required for nutrition and virulence. Interestingly, in the protist human pathogen *Trypanozoma cruzi*, the deletion of the AP-1*γ* encoding gene blocks the transport of the secreted protease cruzipain, a protease involved in nutrition, differentiation and virulence (Moreira et al., 2017). Thus, the clathrin adaptor AP-1 may play a conserved role in the secretion process of enzymes required for nutrition and virulence in eukaryotic pathogens. Intriguingly, other enzymes involved in macromolecule degradation can bypass the AP-1 route: the secretion of several pectin-hydrolases seems to be AP-1 independent. The molecular actors contributing to the AP-1 independent secretion of hydrolytic enzymes remain to be identified in plant-pathogenic fungi. Their discovery would bring a novel understanding of the general secretion mechanisms in Fungi.

### AP-1 is a virulence determinant in plant-pathogenic fungi

The capacity of *Botrytis cinerea* to infect plants depends on the formation of infection cushions (ICs), multicellular appressoria dedicated to plant penetration and secretion of virulence factors (de Vallée et al., 2019; Choquer et al., 2021). The AP-1^cond^ mutant is impaired in the differentiation of these infectious structures, as are *Botrytis* clathrin mutants (Souibgui et al., 2021). Souibgui et al., (2021) suggested that clathrin participates in the differentiation process of IC, possibly through the formation and/or trafficking of vesicles involved in hyphal growth or the transport of IC-specific actors. Our study reveals that the clathrin adaptor AP-1 is required for IC formation, comforting this hypothesis.

The capacity of the necrotroph *B. cinerea* to degrade plant cells is related to the secretion of numerous hydrolytic enzymes (van Kan, 2006). Considering the exoproteome of the AP-1^cond^ mutant, not only several lipases and the pectinase BcPG2 were strongly down-accumulated but also several proteins involved in the degradation of plant polysaccharides and the production of ROS. This defect explains the absence of necrosis observed in the mutant.

Overall AP-1 is involved in many virulence-related biological processes: growth for host-colonization, secretion of hydrolytic enzymes for plant tissue degradation, and establishment of a functional cell-wall that protects the fungus from environmental stresses including plant defenses. In the devastating wheat pathogen *Fusarium graminearum*, the disruption of AP1-σ, another subunit of the complex showed pleiotropic phenotypes including defects in growth, sporulation, mycotoxin production and pathogenicity. The deletion mutant was unable to infect flowering wheat heads and coleoptiles (Wu et al., 2023). In other eukaryotes, AP-1 has also been described as a virulence determinant in protozoan parasites such as *Trypanosoma cruzi* and *Leishamania mexicana mexicana* (Vince et al., 2008; Moreira et al., 2017). Hence, AP-1 might be a conserved and critical virulence determinant in eukaryotic pathogens

## Conclusion

Our work demonstrates that the clathrin adaptor AP-1 is essential for several biological functions such as fungal growth and particularly hyphal polarity, cell-wall formation, and secretion of proteins involved in tissue maceration. These pleiotropic functions of AP-1 highlight the importance of the clathrin/AP-1 machinery in the biology and beyond the infection process of a phytopathogenic fungus . The identification of clathrin/AP-1 secretory cargoes that may be involved in fungal pathogenicity remains a challenge but will provide a new understanding of the secretion processes involved in virulence in pathogenic fungi and other eukaryotic pathogens.

## Abbreviations

TEM: transmission electron microscopy
CAZy: Carbohydrate Active Enzymes
FCWE: Fungal Cell Wall Enzymes
CDIP: Cell Death Inducing Proteins
TM: Transmembrane Domains
SP: Signal Peptides
ER: Endoplasmic Reticulum

## Acknowledgments

The authors are grateful to Mathieu Lays for his contribution to the measurement of secreted enzymes activities.

