## Supporting Information for "Role of the clathrin adaptor AP-1 in cell wall integrity and virulence factor secretion in the plant pathogen Botrytis cinerea"

**Appendix S1**

**Experimental procedures**

**Construction of the *ΔBcap1b,* AP-1$^{cond}$ mutants and the complemented strain AP-1$^{cond}$/C**

Knock-out transformants and knock-in mutants were constructed using a gene replacement strategy. Genetic cassettes were constructed by double-joint PCR (Yu, et al., 2004). All primers used are listed in Table S1.

***ΔBcap1b* strain:**

For *ΔBcap1b* construction (Figure S1A), 5' and 3' flanking regions of *Bcap1b* were amplified by PCR from *Botrytis cinerea* genomic DNA using primer pairs P1/P2 and P3/P4. The hygromycin resistance cassette *hph* (1363 pb) was amplified using primer pair P5/P6. Assembly of fragments was performed by double-joint PCR, and the deletion cassette was amplified with primer pair P7/P8 to form the *ΔBcap1b* cassette. The deletion cassette was then transferred into the P7 vector to produce the p7-*ΔBcap1b* plasmid. The plasmid was verified by sequencing. *Agrobacterium tumefaciens* mediated transformation (*At*MT) was used for the transformation of *Botrytis cinerea* using the protocol described by Roland et al., 2003 Basically, *Agrobacterium tumefaciens* (LBA1126) was transformed with p7-*ΔBcap1b* and cultured with *B. cinerea* conidia on cellophane-covered agar plates for 2 days, 21°C. Then, cellophane sheets were transferred to selective media (Tanaka medium, hygromycin B) for 5-7 days at 21°C.

Diagnostic PCR was performed to confirm that hygromycin-B-resistant transformants carried *hph* and homologous using the primer pair P11/P14. To obtain homokaryotic lines of transformants, several rounds of single-spore isolation on culture medium containing hygromycin-B (70 $\mu g.ml^{-1}$) were performed.

**AP-1 $^{Cond}$ strain:**

For the construction of the *pNiaD::Bcap1b* strain (named AP-1 $^{Cond}$) (Figure S2.A), four DNA fragments were required. The 5' flanking region of *Bcap1b* (1041 bp), the *niaD* promoter (1475 bp), and the 5' end of *Bcap1b* coding sequence (1040 bp) . They were amplified from *Botrytis cinerea* genomic DNA using the primer pairs P1/P2, P21/P22 and P23/P24 respectively. The hygromycin resistance cassette *hph* was amplified using primer pair P3/P20.

The gene replacement cassette *Bcap1b-hph-pNiad::Bcap1b* was obtained with primer pair P29 /P30 and cloned into the P7 vector and verified by sequencing before introduction into the *A. tumefaciens* LBA1126 strain.

Homokaryotic lines of transformants were obtained by several rounds of single-spore isolation on MMIII minimal medium (2 g/L, $NaNO_3$) containing hygromycin-B (70 µg.ml$^{-1}$).

Absence of *Bcap1b* parental locus and detection of *pNiaD::Bcap1b* insertion at the targeted locus was verified by PCR. The primer pairs are indicated in Figure S2.

**AP-1 $^{cond}$/C strain:**

For complementation of the *pNiaD::Bcap1b* mutant (Figure S2.B), the *p7-nptII-pOliC-Bcap1b* vector was constructed. The *nptII* gene confers resistance to Geneticin G418, and *Bcap1b* is under the control of the promoter *pOliC* from *A. nidulans*. The *OliC* promoter and the *Bcap1b* coding sequence and terminator were amplified by PCR and assembled by double-joint PCR to obtain the *pOliC::Bcap1b* cassette that was transferred into the *p7-nptII* vector by IVA cloning (Garcia-Nafria et al, 2016). The plasmid was verified by sequencing.

*A. tumefaciens* transformed with *p7-nptII-pOliC-Bcap1b* vector was used for *At*MT of the ***AP-1*** $^{Cond}$ mutant to produce the complemented strain AP-1 $^{cond}$/C. *B. cinerea* transformants were selected using Geneticin G418 (150 µg.ml$^{-1}$). Diagnostic PCR was performed to confirm that geneticin-resistant transformants carried *nptII*$^{R}$ using primer pair 47/48

**Construction of the BcCHSIIIa-GFP and AP-1 $^{cond}$-BcCHSIIIa-GFP strains:**

GFP fused to the C-terminal end of BcCHSIIIa was introduced in the parental and AP-1$^{cond}$ strains using *At*MT. A gene replacement strategy was developed (Figure S7). The genetic cassette *BcchsIIIa-gfp-tniaD-nat*$^{R}$*-3'UTR-term* was produced by double-joint PCR. *nat*$^{R}$ includes the *nat1* gene placed under the control of the *OliC* promoter and confers resistance to nourseothricin. The 3' end of *bcchs3a* encoding sequence excluding stop codon (1031 bp), the 3'UTR *bcchs3a* and terminator region (924 pb) were amplified from *Botrytis cinerea* genomic DNA (50 ng) using primer pairs P51/P52, and P55/P56, respectively (Table S1). The $_{e}$*gfp-tniaD-nat*$^{R}$ fragment (2494 bp) was amplified from the p7-BcCHSVa-GFP vector using primer pair P53/P54. The *Bcchs3a-*$_{e}$*gfp-tniaD-nat*$^{R}$*-3UTR-term* cassette was amplified with primer pair P57/P58 and transferred to the *p7* vector by IVA cloning. The plasmid was verified by sequencing and introduced into the parental strain and the AP-1$^{cond}$ mutant using *At*MT to produce *BcCHSIIIa-GFP* and AP-1$^{cond}$*pNiaD::Bcap1b-BcCHSIIIa-GFP* strains respectively. *B. cinerea* transformants were selected using nourseothricin (70 µg.ml$^{-1}$).

Homologous recombination was detected by diagnostic PCR on nourseothricin-resistant transformants using primer pairs P61/P62 and P63/P64.

**A**

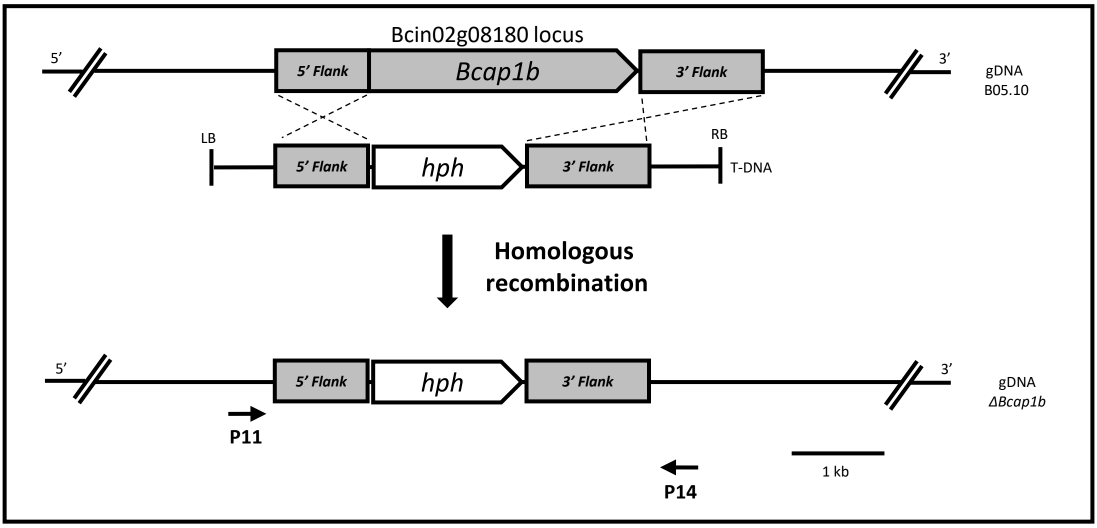


**B**

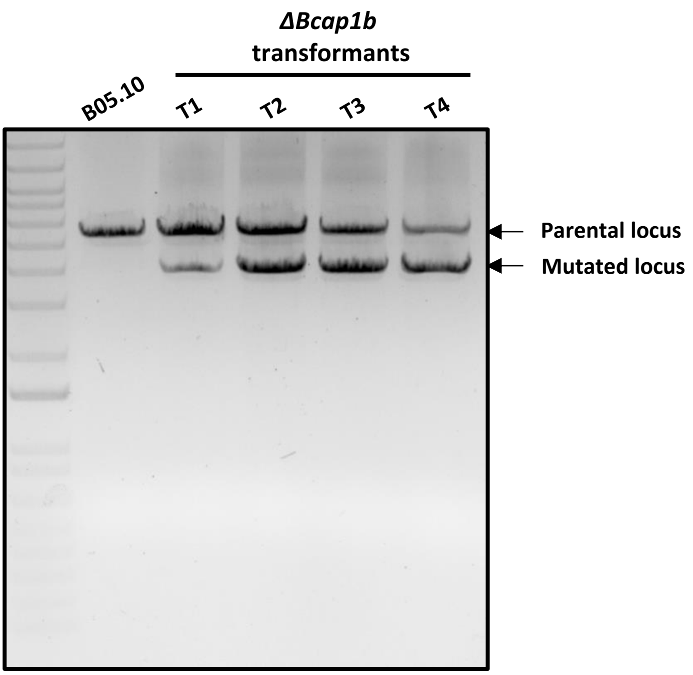


**Figure S1.** Construction and identification of Bcap1b null mutants. **(A)** Scheme of the gene replacement strategy used to create *ΔBcap1b* mutants. Dashed lines: expected double recombination between the Bcap1b 5' and 3' flanking regions present in the targeted locus and in the *A. tumefaciens* T-DNA carrying the selection cassette. **(B)** Diagnostic PCR of *ΔBcap1b* transformants after 5 rounds of single-spore isolation: presence of parental locus was detected using primer pair P11/P14 (parental locus: 5.1 kb; mutated locus: 3.8 kb).

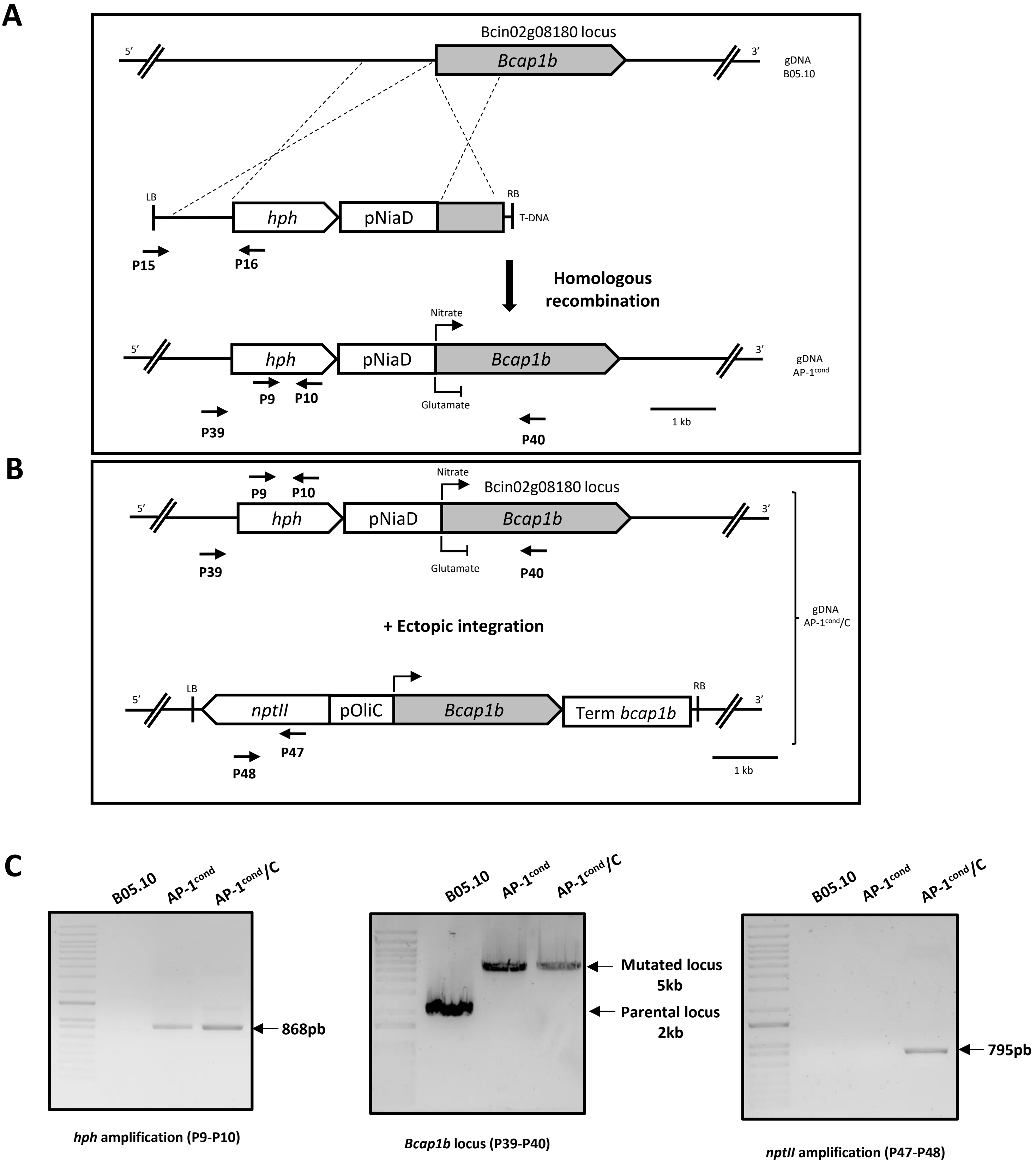


**Figure S2.** Construction and molecular characterization of the pNiaD::bcap1b (AP-1$^{cond}$) and the complemented AP-1$^{cond}$/C strains. **(A)** Scheme of the gene replacement strategy used to create the AP-1$^{cond}$ mutant. Dashed lines: expected double recombination between the Bcap1b 5' and 3' flanking regions present in the targeted locus and in the T-DNA carrying the conditionnal cassette. **(B)** Scheme of ectopic integration of the complementation DNA cassette used to create the AP-1$^{cond}$/C strain. **(C)** Diagnostic PCR of the AP-1$^{cond}$ mutant and the complemented strain AP-1$^{cond}$/C. Primers used are indicated. at the bottom of gel images and as arrows in (A) and (B).

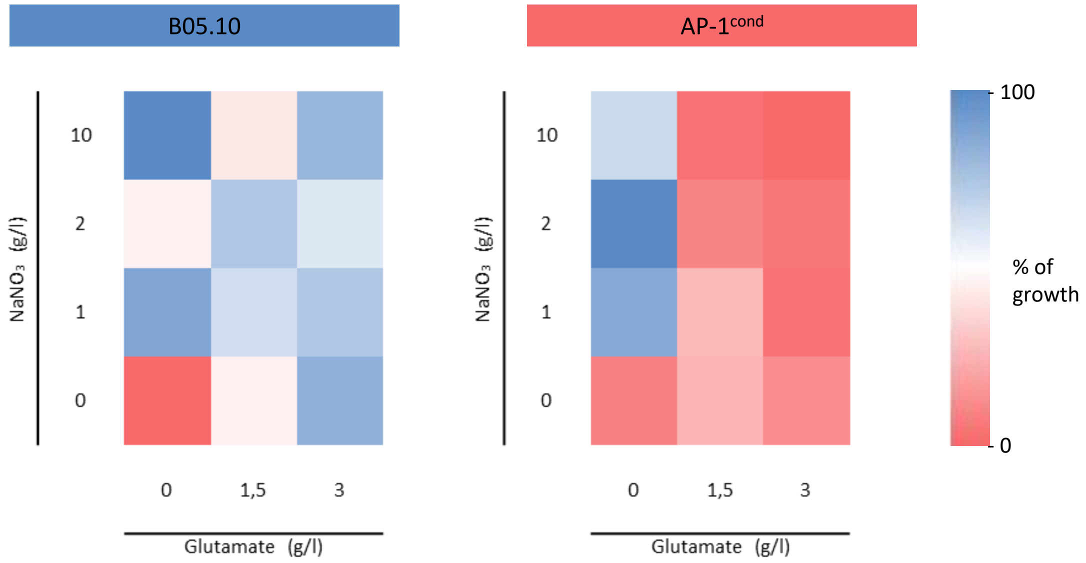


**Figure S3.** Growth of the parental and *AP-1*$^{cond}$ mutant strains in media containing various concentrations of $NaNO_3$ and glutamate. 600 spores per well were inoculated in a 96-well microplate containing media. Percentages of growth were calculated from OD $_{620}$ after 7 days of incubation at 21°C. All experiments were conducted in triplicates with 4 technical replicates per condition.

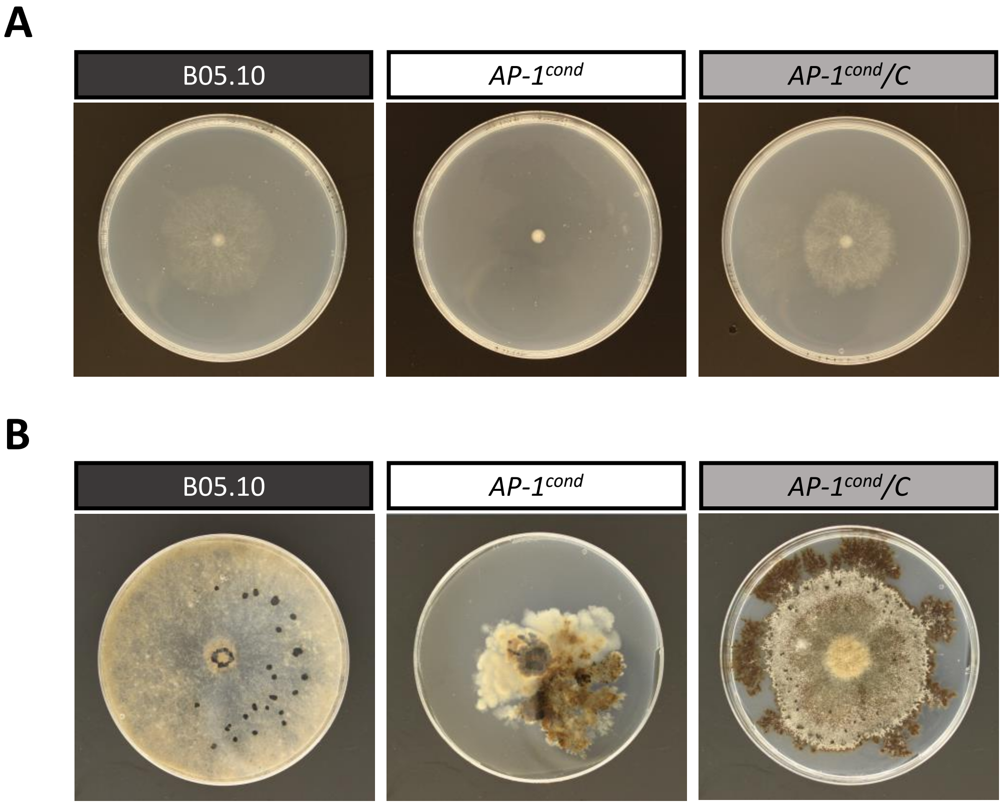


**Figure S4.**. Radial growth of the parental B05.10, *AP-1*$^{cond,}$ *AP-1*$^{cond}$*/C*. **(A)** Strains on solid minimal nitrate medium after 7 days of incubation at 21°C and **(B)** on modified Tanaka medium after 29 days at 21°C.

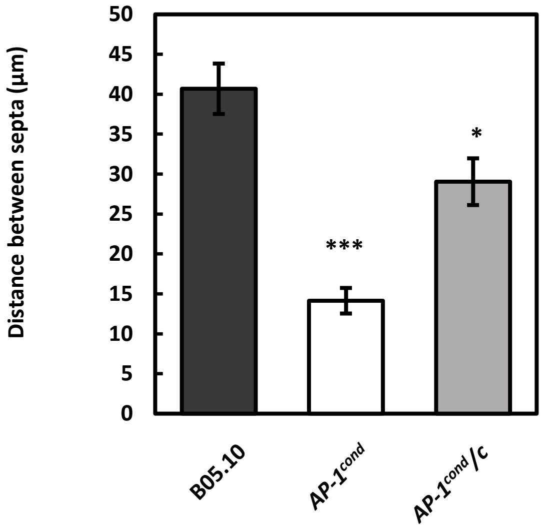


**Figure S5.** Distance between septa in B05.10, AP-$1^{cond}$ and the complemented AP-$1^{cond}$/C strains. Hyphae were observed after 48h of growth in liquid minimal medium supplemented with NaNO3. Three independent cultures were performed and more than 30 distances between septa were measured for each experiment. Means with standard deviations are indicated, asterisks indicate signifuicant difference compared to the B05.10 strain (Student's t-test,* p-value <0.05; *** p-value <0.001).

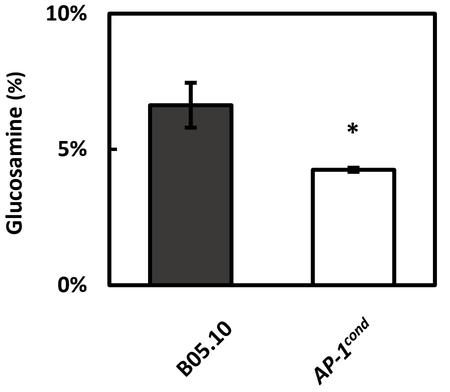


**Figure S6.** Quantification of free N-acetyl glucosamine. Mycelia were grown in in liquid MMII medium containing nitrate for three days under agitation at 21°C. The OD250 were converted to µg of D-glucosamine using 1 $OD_{250}$=8 µg glucosamine. The results are expressed as a % of glucosamine in dry mycelium. Means and standard were calculated from 3 independent experiments (n=9)( Student's t-test, * p-value<0,05)

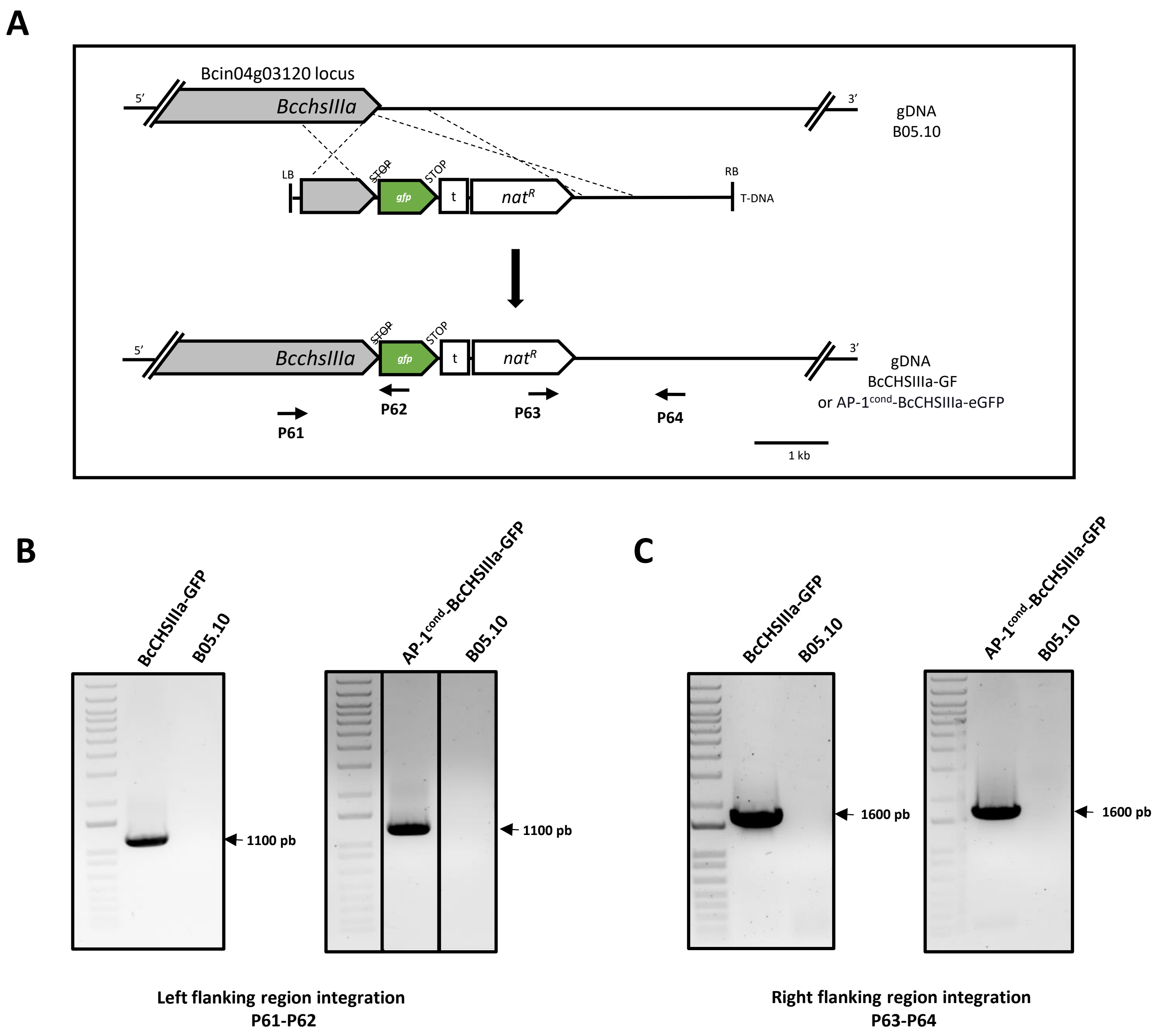


**Figure S7** Construction of the BcCHSIIIa-eGFP and AP-1$^{cond}$-BcCHSIIIa-eGFP strains**. (A)** Scheme of the gene replacement strategy used to create the strains. Dashed lines: expected double recombination between the BcchsIIIa 5' and 3' flanking regions present in the targeted locus and in the *A. tumefaciens* T-DNA carrying the translationnal-fusion cassette. **(B)** Diagnostic PCR of left flanking region integration of BcCHSIIIa-GFP and AP-1$^{cond}$-BcCHSIIIa-eGFP strains **(C)** Diagnostic PCR of right flanking region integration of BcCHSIIIa-GFP and AP-1$^{cond}$-BcCHSIIIa-eGFP strains . Primers used are indicated at the bottom of gel images and as arrows in (A). t= niaD terminator

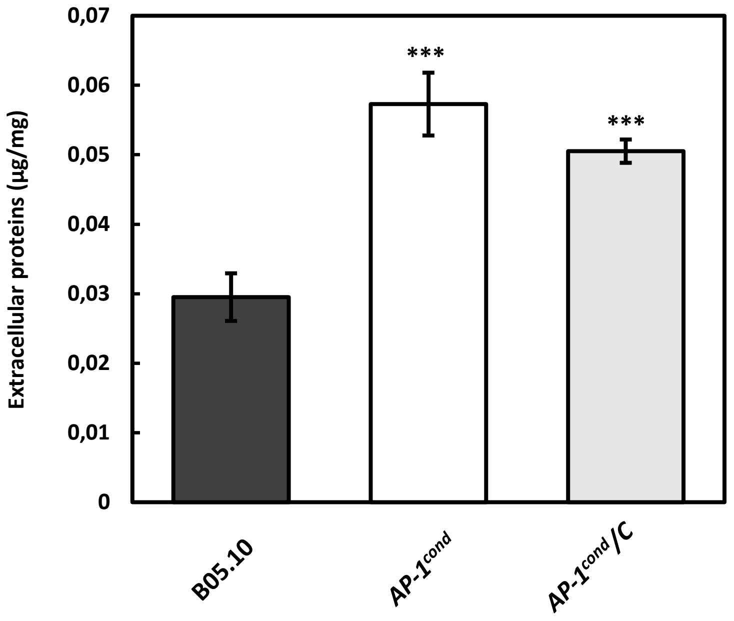


**Figure S8.** Quantification of extracellular proteins in minimal liquid medium (supplemented with NaNO3) after three days of culture. Four independent replicates were performed . Standard deviations are indicated, and asterisks indicate significant difference compared to the parental B05.10 strain (Student's t-test ***<0.001).
